# Updated binding model for a splicing factor, PTBP1 (Polypyrimidine Tract Binding Protein1)

**DOI:** 10.1101/2025.08.08.669377

**Authors:** Areum Han, Junehawk Lee

## Abstract

RNA-binding proteins (RBPs) are central regulators of post-transcriptional gene expression, and identifying their RNA targets is essential for understanding transcriptome dynamics. Predictive computational models enable the identification of RBP binding sites from RNA sequences *in silico*, complementing experimental assays. We previously developed a PTBP1 binding model based on a Hidden Markov Model (HMM) that utilized overlapping nucleotide triplets but was limited due to the assumption of independence between triplets. In this study, we present an improved PTBP1 binding model using Transcription Factor Flexible Models (TFFMs), which relax this assumption and accommodate flexible-length motifs. Trained on PTBP1 CLIP-seq data from HeLa cells, the TFFM-based approach captures positional dependencies among nucleotides, consistent with known PTBP1 binding motifs, and achieves improved predictive performance. The updated model shows a stronger correlation with experimental binding affinities (r = −0.95) compared to the previous HMM (r = −0.91) and maintains specificity by correctly scoring non-binding RNA substrates. This work highlights the advantage of flexible, context-aware modeling frameworks in predicting RBP–RNA interactions and offers a more accurate tool for studying PTBP1 binding specificity.

## INTRODUCTION

Mammalian cells have several hundred RNA-binding proteins (RBPs). RBPs are the main conductors of post-transcriptional gene expression programs. Identifying their RNA targets is crucial to understanding the mechanisms underlying transcriptome dynamics. Ideally, we would be able to predict the binding affinity of RBPs to any RNA solely by scanning its nucleotide sequence. Computational models have been developed that allow us to scan the genome and identify targets *in silico*. These models often help identify novel targets in low-mappability, repeat-rich regions that could be missed by binding assays followed by sequencing, such as CLIP-seq.

In previous study, we built a binding model for PTBP1 based on a statistical framework, a Hidden Markov Model (HMM). The predicted binding scores from this model were highly correlated (r□= □−0.91) with experimentally determined dissociation constants (Kd values).

The model is based on triplets (three nucleotides) to assess whether triplets would segregate into two states and whether these two states differ in their PTBP1 binding or non-binding potential. In the previous model, triplets were sampled from the subject RNA sequence in a sliding-window fashion, allowing overlaps. Although the performance is good, the overlaps between neighboring triplets could not completely satisfy the HMM assumption that the current observation is independent of the previous observation.

We tried to tackle this issue by adopting how a gene prediction program approaches codons, but the performance was not good. In detail, we sampled non-overlapping triplets with three different starting positions and chose the best-scored triplet set. Probably, the above scoring did not perform well because it was not able to model the flexible length of binding and non-binding states. It is hard to imagine that the lengths of binding and non-binding regions are exactly multiples of three.

Two key binding properties of PTBP1 are: 1) the lengths of nucleotide regions corresponding to binding and non-binding states are flexible, and 2) there is positional interdependence within short elements. A new statistical framework, Transcription Factor Flexible Models (TFFMs), was developed to relax the independence assumption while accommodating flexible-length motifs for transcription factors [2]. We updated the PTBP1 binding model using the TFFM framework. The updated model not only corrected the mathematical flaw from the previous model but also improved predictive performance.

## METHODS AND RESULTS

The HMM is composed of a set of hidden states emitting nucleotides with a corresponding set of emission probabilities, and a set of transition probabilities from state to state. These probabilities can be trained using high-throughput data. In this study, we used the same training set that we used in our previous paper [1]: PTBP1 CLIP-seq data in HeLa cells. This dataset is composed of sequences of CLIP clusters from CLIP tags, extended by 50 nucleotides on both sides, from both PTBP1 monomer and dimer ribonucleoprotein complexes.

For initialization, first we removed short sequences (<9 nt) and randomly sampled 10% of sequences from each chromosome. Using MEME software, we derived the most overrepresented motifs. Because the TFFM framework was originally designed for training DNA-binding transcription factors, we replaced uracil with thymine in the MEME output for compatibility. We trained TFFMs after initializing them using the MEME output.

Figure 1 describes the 1st-order TFFM model trained for the PTBP1 binding motif. Figure 1A and B are sequence logos representing the PTBP1 binding sites. As depicted in Figure 1A, the probabilities of a nucleotide appearing at a certain position of the motif are represented based on the nucleotide at the previous position. Interestingly, the transition from position 1 to 2 and 2 to 3 prefers T (T: uridine in RNA) to C and C to T transitions, respectively, but T to T is preferred in the transition from position 3 to 4. The UCUU sequence (shown here as TCTT) has been a highly significant word in previous reports and consistently appears in the summary logo again, as shown in Figure 1B. Therefore, TFFM allowed us to model the RNA binding motif by incorporating sequence context and describing it more precisely. The detailed description of the trained 1st-order TFFM for the PTBP1 binding motif is shown as an HMM schema in Figure 1C.

**Figure 1.**
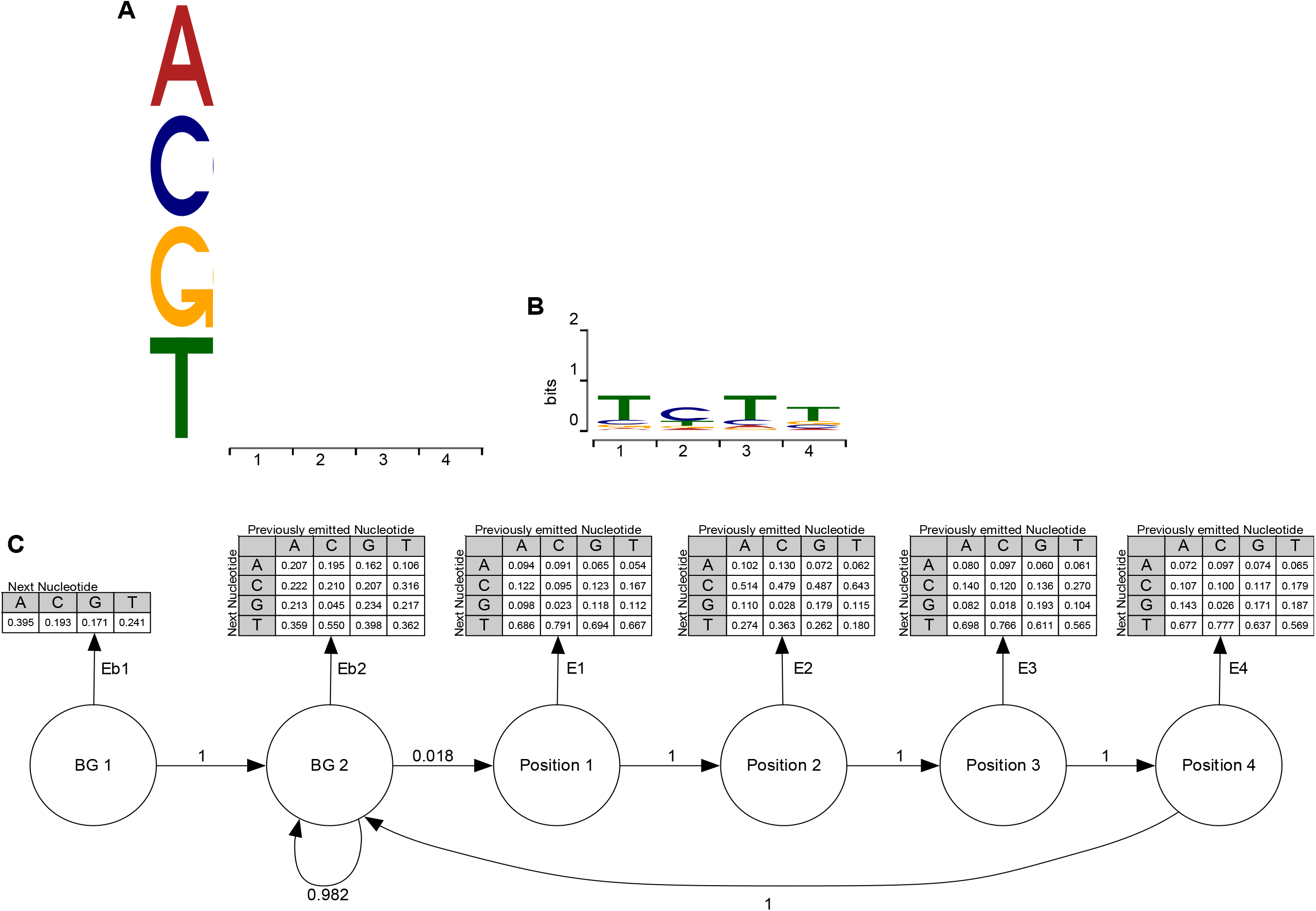
Trained 1st-order TFFM model for the PTBP1 binding motif. (A) Dense logo of the model (A: adenine, C: cytosine, G: guanine, and T: uracil in RNA). Each column represents a position within the PTBP1 binding motif, and each row corresponds to the probabilities of a nucleotide appearing based on the nucleotide at the previous position. Higher opacity indicates a higher probability of observing the nucleotide at that position, depending on the nucleotide at the previous position. (B) Summary logo of the model. (C) HMM schema of the model. The first two states represent the background (BG), and the other four states denote consecutive positions within the PTBP1 motif. The emission probabilities of each state are shown in table form, illustrating the probability of each nucleotide emission depending on the previously emitted nucleotide.

To evaluate the trained TFFMs, we scanned the PTBP1 binding sites, calculated the probability of occupancy (Pocc) scores, and checked whether the Pocc score was well correlated with the measured binding affinity of PTBP1. We ignored the reverse strand match, as we were scoring single-stranded RNA binding events.

As shown in Figure 2, the Pocc of the 1st-order model is indeed more strongly correlated (r = −0.95) with the actual RNA-binding affinity than the previously published HMM model (r = −0.91). Next, we tested whether the new model could also distinguish non-PTBP1 targets, or negative targets. Previously, RNA substrates 1, 2, 3, and 5 did not bind PTBP1 within the tested protein concentration range and received negative scores from the earlier model. As shown in the table, these RNA substrates were again scored low by the new model (Pocc < 0.4).

**Figure 2.**
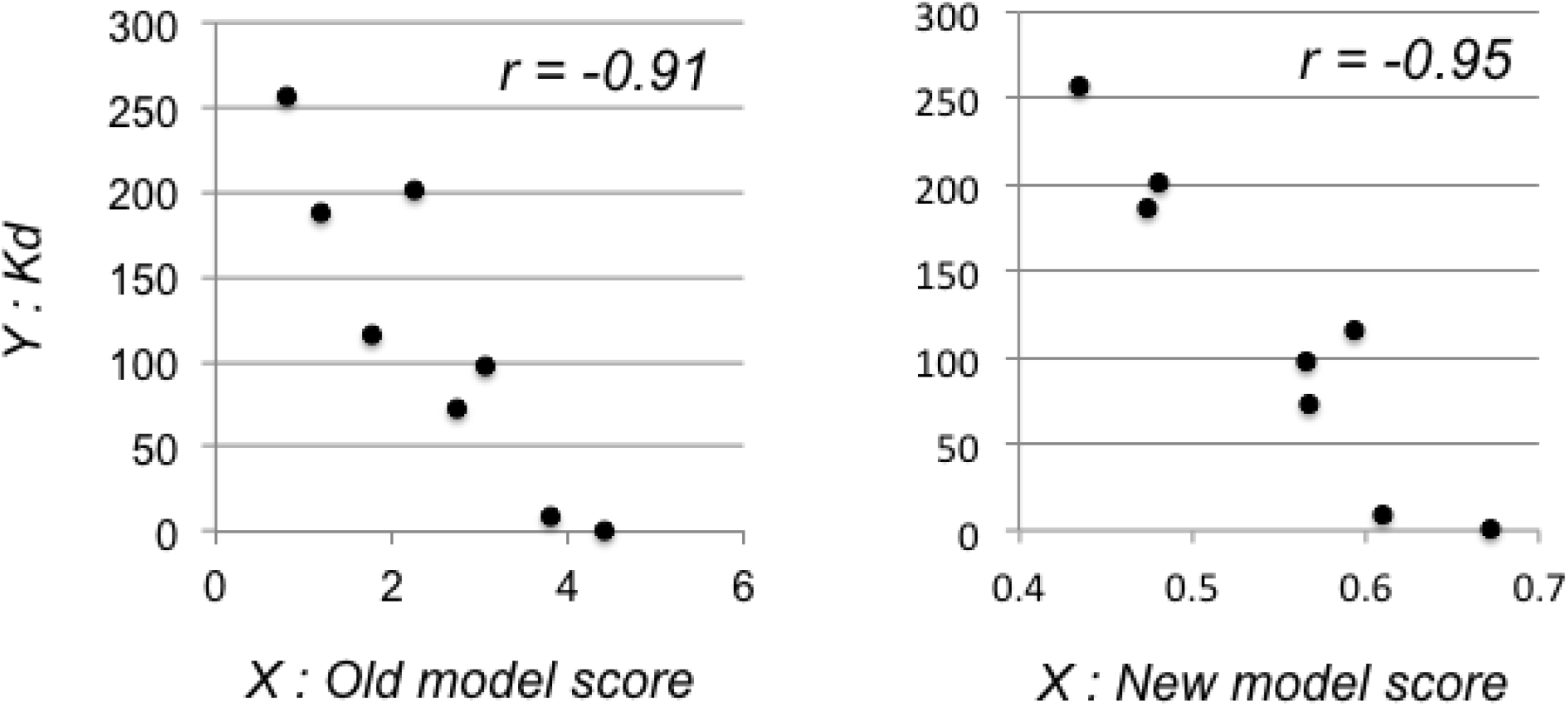
The new model based on the first-order TFFM showed better performance than the previous model in predicting PTBP1 binding affinity (r = Pearson correlation).

**Table 1.**
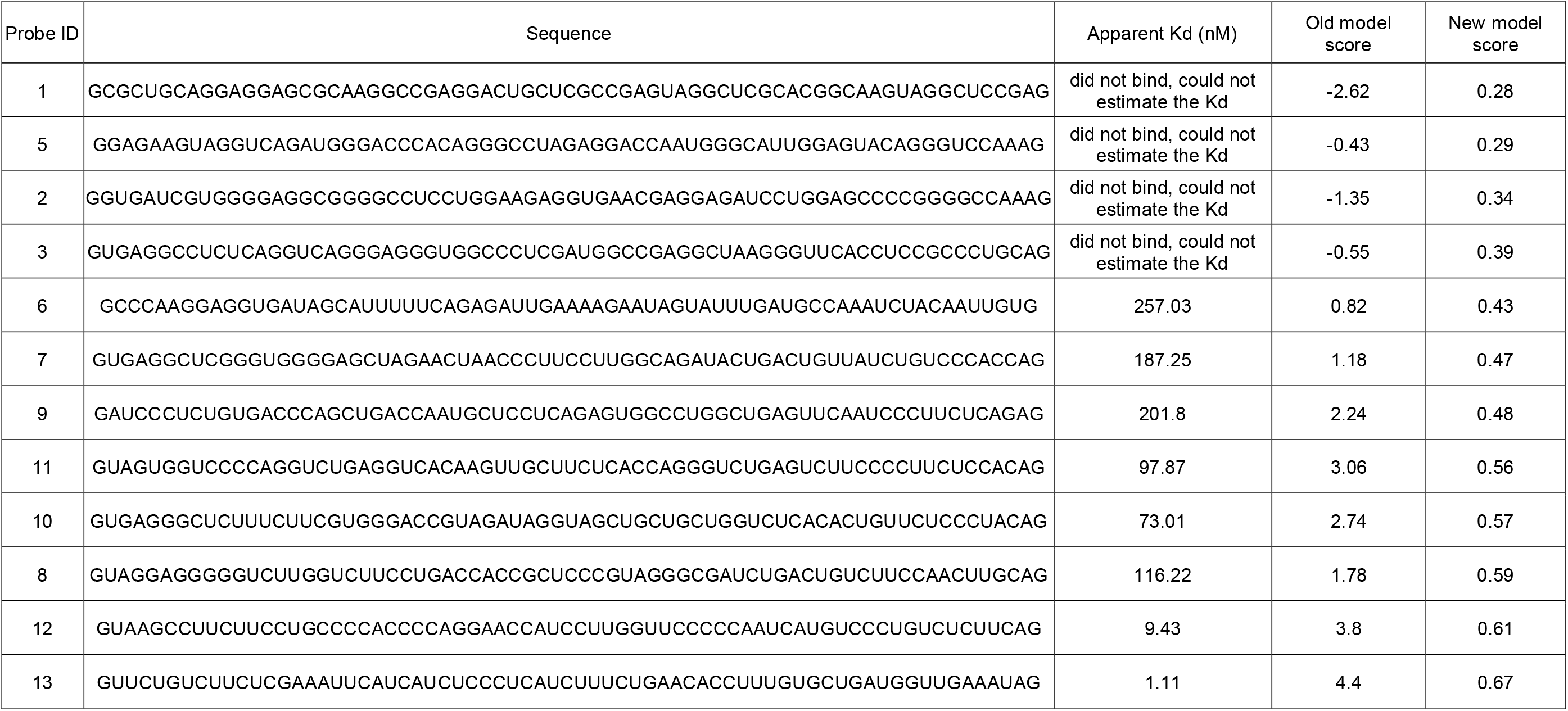
RNA substrates and corresponding Kd values and scores (sorted by the new model score).

## ACKNOWLEDGMENT

Authors thank Yaron Orenstein, who initially introduced the TFFM framework.

